# Gene regulatory networks associated with lateral root and nodule development in soybean

**DOI:** 10.1101/2019.12.15.876995

**Authors:** Shuchi Smita, Jason Kiehne, Sajag Adhikari, Erliang Zeng, Qin Ma, Senthil Subramanian

## Abstract

Legume plants such as soybean produce two major types of root lateral organs, lateral roots and root nodules. A robust computational framework was developed to predict potential gene regulatory networks (GRNs) associated with root lateral organ development in soybean. A genome-scale expression dataset was obtained from soybean root nodules and lateral roots and subjected to biclustering using QUBIC. Biclusters (BCs) and transcription factor (TF) genes with enriched expression in lateral root tissues were converged using different network inference algorithms to predict high confident regulatory modules that are repeatedly retrieved in different methods. The ranked combination of results from all different network inference algorithms into one ensemble solution identified 21 GRN modules of 182 co-regulated genes networks potentially involved in root lateral organ development stages in soybean. The pipeline correctly predicted previously known nodule- and LR-associated TFs including the expected hierarchical relationships. The results revealed high scorer AP2, GRF5, and C3H co-regulated GRN modules during early nodule development; and GRAS, LBD41, and ARR18 co-regulated GRN modules late during nodule maturation. Knowledge from this work supported by experimental validation in the future is expected to help determine key gene targets for biotechnological strategies to optimize nodule formation and enhance nitrogen fixation.

## Introduction

Gene regulation is a fundamental process that controls spatial and temporal patterns of gene expression. Transcription factors (TFs) are central to gene regulation as their activities determine the expression patterns and function of multiple genes (1). A TF is a functional protein that binds to short sequences (called TF binding site; TFBS or *cis*-regulatory elements) on the upstream promoter region of genes to regulate their transcription. One TF can regulate multiple genes including other TFs in a signaling, developmental, or metabolic pathway and so act as master regulators of the pathways. The nested group of all different TF regulators and their downstream target genes form gene regulatory networks (GRNs) (2). Identification of gene regulatory networks and key TFs that are part of these networks is an effective approach to answer multiple biological questions on genotype to phenotype relationships. For example, potential TFs, their co-regulators, and downstream signaling pathways, and target genes associated with specific biological processes can be predicted by constructing GRNs.

Clustering of large-scale datasets such as global gene expression profiles obtained by RNA-sequencing to identify co-regulated TFs and the targeting genes is a promising approach to model and infer the GRNs at a systems level (3, 4). Briefly, genes/TFs with similar expression patterns (i.e. co-expressed genes) with a tendency to co-activate across a group of samples might give insight on TFs regulated gene network and related biological process. In fact, multiple levels of gene regulation affect transcriptional regulatory capabilities (5). Recruitment and binding of other protein such as “co-factors” in complexes and other small protein molecules to target DNA sequences is one of the major mechanisms (6). Often, this interactions between different TFs and co-factor partners are studied using protein-protein interaction (PPI) assays which provide immediate insights into their potential biological function (7, 8). GRNs can be validated by PPI data, as PPIs can reveal signaling, regulatory and/or biochemical roles of proteins based on their interactomes (9).

The combined use of high-throughput data and mathematical models to build gene co-expression and regulatory networks is the core principle of systems biology approaches (10). However, these large-scale datasets are likely to be noisy, and GRN predictions using these big datasets may contain many false positives. Additionally, GRN inference is a computationally intensive job; so filtered datasets consisting of well-defined/accurate datasets (such as significantly co-expressed genes set) might dramatically reduce the computational complexity and time. Most importantly, it would reduce the true search space for the prediction of regulators (TFs) and their potential target genes. In order to obtain significantly co-expressed genes, “biclustering” is a desirable method as it allows two-way clustering of genes as well as samples i.e. a similar expression pattern (co-expressed genes) under a subset of all samples. Subsequently, this sorted biclustering-filtered data fed into GRN inference algorithms might improve and accurate predictions of a regulator and their target genes. We applied this approach to determine gene regulatory networks associated with root lateral organ development in soybean.

Plants produce lateral organs such as leaves, flowers, and axillary branches in the shoot, and lateral roots in the roots. Pools of stem cells present in the growing tip of the shoot (the shoot apical meristem) contribute to the formation of aerial/shoot lateral organs. Lateral organs in the root are unique in that they are derived via “*de novo*” differentiation of mature cells in the root. Lateral roots are present in all vascular plants, but a group of *Fabids* clade plants is capable of producing another root lateral organ, called root nodules. These arise from specific and coordinated interactions with a set of nitrogen-fixing bacteria collectively called rhizobia. For example, the interaction of soybeans with *Bradyrhizobium diazoefficiens* results in root nodules. Biological nitrogen fixation in root nodules helps reduce the need for chemical nitrogen fertilizers, which are expensive and cause environmental pollution. Similarly, proper patterns of lateral root formation (root branching) are crucial for plants to access water and other nutrients in the soil. Therefore, these two root lateral organs play important roles in the development of soybeans, a major crop in the United States as well as in other countries. Many functional genomics studies have identified genes expressed during nodule development in soybean and other legumes, but gene expression profiles during lateral root formation have not been evaluated in legumes (11, 12).

Recently, we obtained transcriptomes of emerging nodules, mature nodules, emerging lateral roots, and young lateral roots in soybean (13), we present a robust computational framework, which we applied to predict TFs and their target GRNs associated with soybean root nodule development. This approach consists of the following steps (Figure 1): (i) preparing a compendium of soybean lateral organ transcriptome data and cataloging TFs enriched in root nodules; (ii) Initial biclustering of transcriptome data using QUBIC (14, 15, 16) to identify all (nodule development stage-specific) co-expressed gene modules; (iii) GRN construction and inference based on identified gene modules and reliable network construction programs, Lemon-Tree (17) and Inferelator (18); (iv) Augmentation of GRNs with evidence from physical or direct and indirect regulatory interaction information from PPI and *cis-*regulatory element enrichment analysis; and (v) building a consensus from different modes of GRN inference for potential regulators and their predicted GRNs. We ran two modes of Lemon-Tree, one with default mode, where Lemon-Tree itself produce the co-expressed clusters and the other mode with reinforced bicluster (BC) information from QUBIC. This study provides a template framework for GRN construction and augmentation by exploiting big data sets, which are increasingly generated, deposited and available (making use of available data) in public domain.

**Figure 1.**
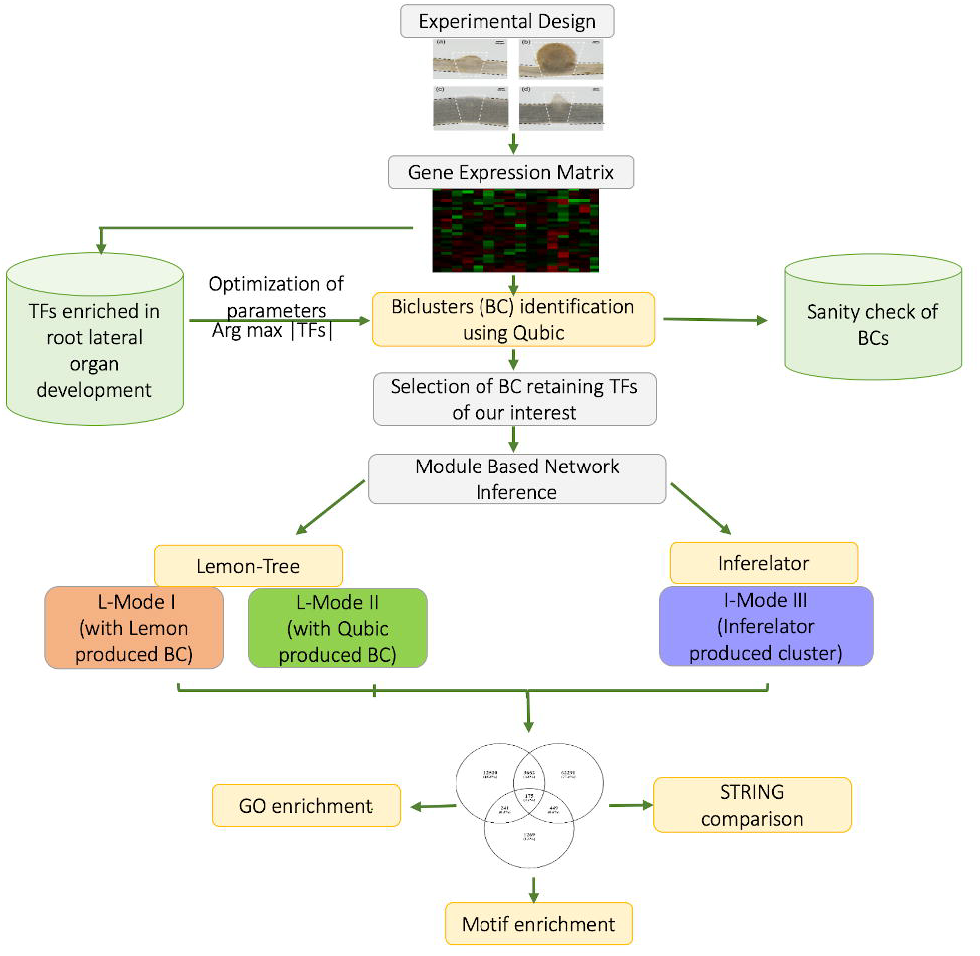
Schematic representation showing our workflows for prediction of regulator transcription factors (TFs) and their Gene Regulatory Networks (GRNs) for root lateral organ development in soybean.

## Material and methods

### RNA-seq dataset for root lateral organ development in soybean

We utilized the genome-wide soybean transcriptome dataset generated for root lateral organs (13). This dataset contains the transcriptomes of two different developmental stages of two root lateral organs collected in three biological replicates: emerging nodules (EN), mature nodules (MN), emerging lateral roots (ELR) and young lateral roots (YLR). Adjacent root sections above and below these organs devoid of any lateral organs (designated as ABEN, ABMN, ABELR, and ABYLR respectively) were used to construct respective age- and inoculation-status appropriate control tissue libraries. Comparison of gene expression profiles between each lateral organ tissue type and the corresponding control tissue type (e.g., EN vs. ABEN, ELR vs. ABELR and so on) helped identify organ-specific/enriched genes. In total, 24 RNA-seq libraries (four target tissue types, four control tissue-types, three biological replicates each) were prepared, sequenced, and analyzed. Expression patterns of preciously known marker genes, consistency between replicates, high sequence quality of this dataset indicated that it was of very high quality and well-suited for global gene expression analysis (13). A total of 113,210 gene transcripts (FPKM threshold ≥ 1 in at least one sample) with their normalized expression values in 24 different tissues from the above dataset were utilized here.

Further, for expression comparisons at different steps during our analysis, we utilized the public datasets, soybean gene atlas encompassing RNA-seq data from 14 different soybean tissues (19) and Soybean eFP browser http://bar.utoronto.ca/efpsoybean/cgi-bin/efpWeb.cgi comprising RNA-seq data from soybean root hair and other tissues (20, 21). Soybean genome sequence assembly version 7.0 (Gmax_109_gene.gff3.gz”; ftp://ftp.jgi-psf.org/pub/compgen/phytozome/v9.0/Gmax/annotation/) was used for gene annotation and Arabidopsis orthologs information.

### Cataloging TFs enriched in root lateral organ development stages in soybean

To achieve our objective of identifying regulator TFs and prediction of GRNs associated with root nodules, we used soybean transcription factor annotations from the Plant transcription factor database (PlantTFDB v3.0; http://planttfdb.cbi.pku.edu.cn/) (22) as a starting point. Among 58 TF families annotated in soybean, 48 TF families had at least one member differentially expressed in at least one of the four organ tissue types. For each TF family, we summed the unique transcripts that were enriched in EN and/or MN to calculate the total number of family members enriched in nodule tissues. Similarly, we calculated the number of TFs enriched in lateral root tissues. By comparing the number of family members enriched in nodule vs. lateral root tissues, we identified nodule-specific or -enriched, lateral root-specific or - enriched, and lateral organ non-specific (equal number of transcripts in lateral root and nodules) TF families (Figure 1; Table S1). Statistical analysis (Fisher’s Exact test, P<0.05) of nodule- vs. lateral root-specific enrichment showed that TALE, MYB-related, MIKC, C2H2, bZIP, G2-like, WRKY, and NFYB were either nodule-specific or significantly enriched in nodules (Figure 2). Overall, very distinct families of TFs appear to be active in nodule and lateral roots despite reported morphological similarities between these organs.

**Figure 2.**
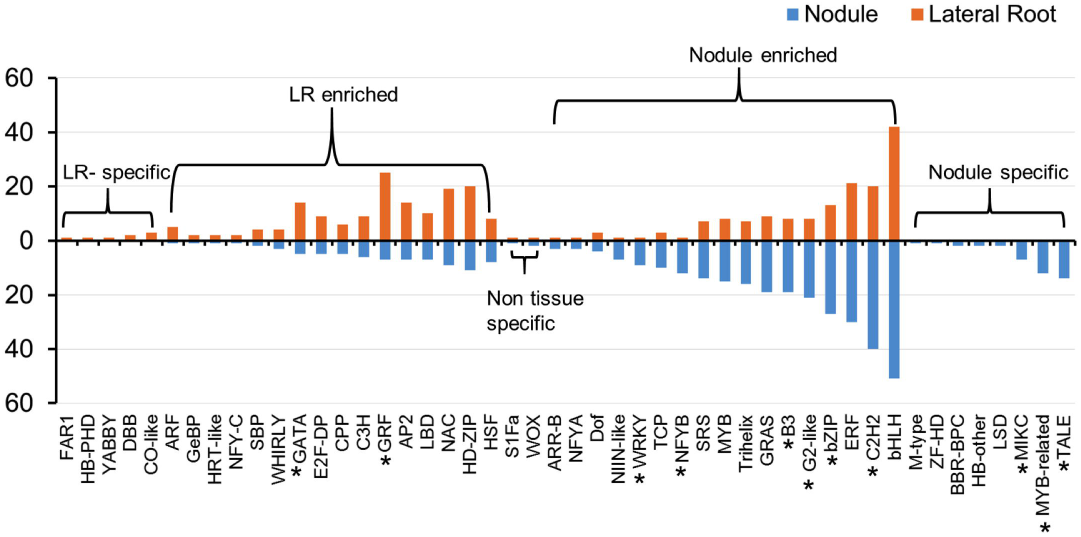
Transcription factor (TF) families enriched in specific root lateral organs. Bar graphs indicated the number of family members enriched in nodules (blue) or lateral roots (orange). TF annotations are based on Plant Transcription Factor Databases (PlantTFDB). Asterisks indicate TF families that were significantly enriched either in nodule or lateral root (Fisher’s exact test; P < 0.05).

We selected a set of 294 TFs, which were differentially expressed and specifically enriched in EN, and MN tissues in our dataset as possible regulators (see Results, Supplementary Table S1). This approach led us to focus on regulators and their GRNs acting specifically during nodule development. We also included 22 previously characterized TFs/ regulator genes reported elsewhere in literature for their respective role in root lateral organ development in model crop plants as positive control marker genes for validation and relevancy of parameters (Supplementary Table S2). For example, ENOD40, FWL1, LBC_A, LBC_C1, LBC_C2, and LBC_C3 genes were used as marker genes, and NIN1 and NSP1 were used as marker regulators for nodule development. ARF5, CRF2, GATA23, LRP1, and TMO7 genes were used as marker regulators for lateral root development. Together, we used 316 TFs of interest as a starting point for the identification of GRNs.

### Initial biclustering of transcriptome data

We utilized normalized expression values of all the 113,210 gene transcripts in 24 libraries for initial biclustering, rather than only significantly differentially expressed gene (DEGs) transcripts. We reasoned that irrespective of enrichment, the TFs and their target gene clusters tend to have similar expression patterns in the root lateral organs, making this an unbiased approach. We chose biclustering (two-way clustering), over traditional clustering for simultaneously clustering using QUBIC (QUalitative Biclustering) (15) to identify all the statistically significant biclusters (BCs) of target genes with TFs, if any as well as samples from the above transcriptome data. Different combinations of QUBIC’s parameters were tuned to optimize biclustering to retain the majority of TFs while keeping the total number of transcripts to the minimum. The program first discretizes the data using the parameters *q* and *r* and then a heuristic algorithm applied to identify biclusters, where *q* is the proportion of affected expression data under all conditions for each gene; and *r* represents the rank of the regulating conditions detected by the parameter *q*. It is suggested to select a smaller *q* to focus on a local regulator (15). Parameter *f* controls the overlap between different BCs, and *k* controls the minimum number of samples in BCs. Another important parameter *c;* which controls the level of consistency in BCs, was tested to balance the number of TFs and a total number of genes covered in BCs. We obtained 219 BCs that contained 240 of the 316 TFs (76%) and 30, 639 out of 113,210 transcripts (~27%; See Results for details). This “filtered” dataset was used for regulator and GRN prediction. All programs were tested and implemented on a Linux server with Intel x86-64 processor and 32 cores with 1TB RAM configuration.

### Prediction of potential TF regulators and their GRN inference

To improve the confidence of regulator and GRN prediction, we utilized two module-based GRN inference methods: Lemon-Tree (v.3.0) (17) and Inferelator (v.2015.08.05) (23). We compared and scored the regulatory prediction made by both methods to select high confidence regulators and their target genes in GRN.

### Lemon-Tree

Lemon-Tree has the option to integrate cluster information; hence, we ran it in two modes: (i) where clusters were generated by Lemon-Tree from the “filtered” dataset (mode I) and (ii) where BC information from QUBIC was fed to Lemon-Tree as co-expressed gene modules for GRN inference (mode II). For mode I, we ran ten independent Gibbs sampler runs of Lemon-Tree (with default parameters) to identify the most confident regulatory modules and TF regulators. The results were used to extract representative module solution (tight clusters) from an ensemble of all possible statistical models using the Gibbs sampler method. Lemon-Tree modules are clustered (hierarchical tree) based on samples with similar mean and standard deviation. This tight cluster corresponds to sets of genes, frequently associated across all clustering solutions. For mode II, we prepared this tight cluster dataset using BCs information from QUBIC, but otherwise used the same settings used for mode I.

In the next step, the Lemon-Tree algorithm provides a list of weighted TFs with a ranked probability score, and the top 1% were selected as true regulators for each cluster of co-expressed genes. A global score reflecting the statistical confidence of the regulator assigned to each node in a hierarchal tree manner for each set of co-expressed genes modules. The regulator score takes into account the number of trees a regulator is assigned to, with what score (posterior probability), and at which level of the tree (24). An empirical distribution of scores for randomly assigned regulators-to-module is also provided to assess significance (17). In this dataset, the lowest score of a regulator in the top 1% list was at least 3x higher than that of the highest score for a randomly assigned regulator (See Result section for details). Therefore, either the top 1% or at least a 3-fold higher score than randomly assigned regulators appears to be a good threshold to determine true regulators.

### Inferelator

Inferelator (20 bootstraps) with default settings was utilized to build regulatory networks. Similar to Lemon-Tree, it also uses the gene expression matrix to predict the regulator TFs and their target genes. However, unlike Lemon-Tree, Inferelator does not take cluster information as input, but generates its own clusters. The program generated a ranked list of target genes for each regulator TF utilizing the gene expression matrix and the TFs of our interest. Unlike Lemon-Tree, there is no "score-based" selection of TFs in Inferelator, while there are score-based regulatory interactions between TF and their target genes. Inferelator-generated scores (s) for TF (*x*) regulating gene (*g)* using input gene expression matrix (*RNA-seq*) as:

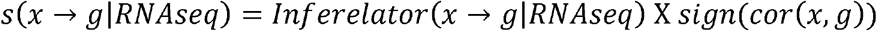

where a regulatory interaction confidence score is multiplied by the sign of the correlation coefficient between the TF and the putative target gene to differentiate putative activating from repressing interactions (positive and negative scores, respectively) (18, 25).

### Combined scoring of regulatory predictions for consensus GRN

By taking advantage of the top regulator prediction feature of Lemon-Tree and top-ranked regulatory target prediction of Inferelator, we compared and combined TF and targeted module genes from all three-inference solutions: Lemon-Tree mode I, II, and Inferelator (described above). The regulatory TFs and corresponding target genes common among all three-inference solutions using Linux “*comm*” command, were rated as potential consensus regulators and their targeted GRN interactions. Ranked score function for every predicted regulatory interaction was calculated by normalizing scores produced by each inference solution (score divided by the highest score in each inference solution) and then averaging normalized score calculated from all three-inference solutions. These ranked scores were used to select high confidence candidate TF-target interactions. These were showed in edges in the GRN modules, visualized and analyzed using Cytoscape (version 3.3.0) (26).

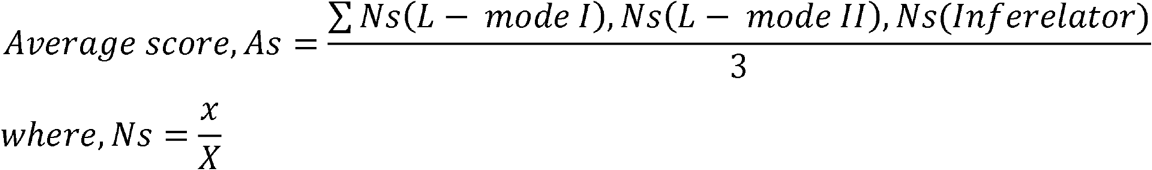

Ns=normalized score

x = probabilistic score from each mode

X= maximum score in each mode

L-mode I = Lemon-Tree mode I

L-mode II = Lemon-Tree mode II

### Protein-protein interaction (PPI) network evidence for physical interaction

Most eukaryotic TFs recruit various co-factors for their DNA-binding specificities and regulatory activities through PPIs. To evaluate potential PPIs that are part of the predicted GRNs, a total of 31,932,066 predicted/experimentally validated soybean protein interactions (NCBI taxon-Id:3847) were obtained from the STRING database (version 10.0) (search tool for the PPI network) (27). This database provides information on both experimental and predicted interactions from varied sources based on co-expression, experiments and literature mining, etc. We evaluated and compared if the predicted TFs and targets from the different inference solutions (Lemon-Tree mode I, II and Inferelator) were potential PPI partners using all the 31,932,066 STRING PPI interactions in soybean. Non-redundant dataset, ignoring the transcript numbers of TFs, targets (from TF-target interactions) predicted by three individual inference solutions and PPI from STRING were compared using the Linux “*comm*” command to identify TF-target pair common in STRING dataset and their PPI scores.

### *Cis*-regulatory motif and functional enrichment analysis evidence for direct regulation

*Cis-regulatory* motif enrichment was carried out using potential promoter sequences of target genes for all potential regulator TFs predicted by all three inference solutions (Lemon-Tree mode I, II and Inferelator). Motif enrichment and Gene Ontology were performed by ShinyGO (http://www.ge-lab.org:3838/go/) using p-value cutoff (FDR) < 0.05 to determine regulation and function.

## Results

### Optimization of QUBIC parameters for initial biclustering

The primary goal for biclustering in our analysis was to optimize the total number of significant BCs; where the majority of the TFs (out of TFs of interest and marker TFs) are retained while keeping the total number of genes to a minimum for true GRN prediction. In order to evaluate this condition, we iterated various runs in several steps to empirically optimize key QUBIC parameters. For example - *q* to focus on a local regulator, and as regulatory networks are quite small networks, we chose smaller *q* values. To control the overlap by checking the overlapping genes and the number of TFs in between produced BCs, we iterated the run with *f =* 0.5 to 0.65 (by 0.1). We used k = 6 presumably to retain at least three replicates each from either early or late developmental stages or from lateral root or nodule tissue types in one BC. Importantly, the consistency level of BCs was tested using parameter “*c*” iterated from *c* = 0.5 to 1 (by 0.1) to balance the number of TFs and a total number of genes covered in BCs. We noticed that the lower consistency level “*c”* values led to the increased size of BCs. We evaluated the produced BCs to determine the *“c”* value at which we covered the greatest number of TFs in comparison to a total number of genes without losing much consistency (*c*). At *c* = 0.98, 76% of the TFs of interest were retained with just 27% of the genes covered in BCs (Figure 3). Interestingly, the maximum number of marker TFs (18 out of 22) cataloged for root lateral organs were covered at *c* = 0.98. On the other hand, at the best consistency level (c=1), only three marker TFs were covered in BCs (not shown). Overall, based on results from several iterations and optimizing for the inclusion of greater number of TFs in BCs, we finalized the following parameters: *r* = 1, *q*= 0.2, *c* = 0.98, *o* = 500, *f* = 0.25, *k* = 6; which produced 219 statistically significant BCs (Supplementary Table S3). These 219 BCs comprised ~27% (30, 639 out of 113,210) of total gene transcripts. Notably, ~76% (240 out of 316 TFs of our interest) of the TFs of interest and marker TFs were retained in 141 of the 219 total BCs produced. The first cluster was the largest cluster with a total of 446 genes. We conclude that the empirical determination of biclustering parameters depending on the biological question and the associated experimental objective is crucial for useful outcomes.

**Figure 3.**
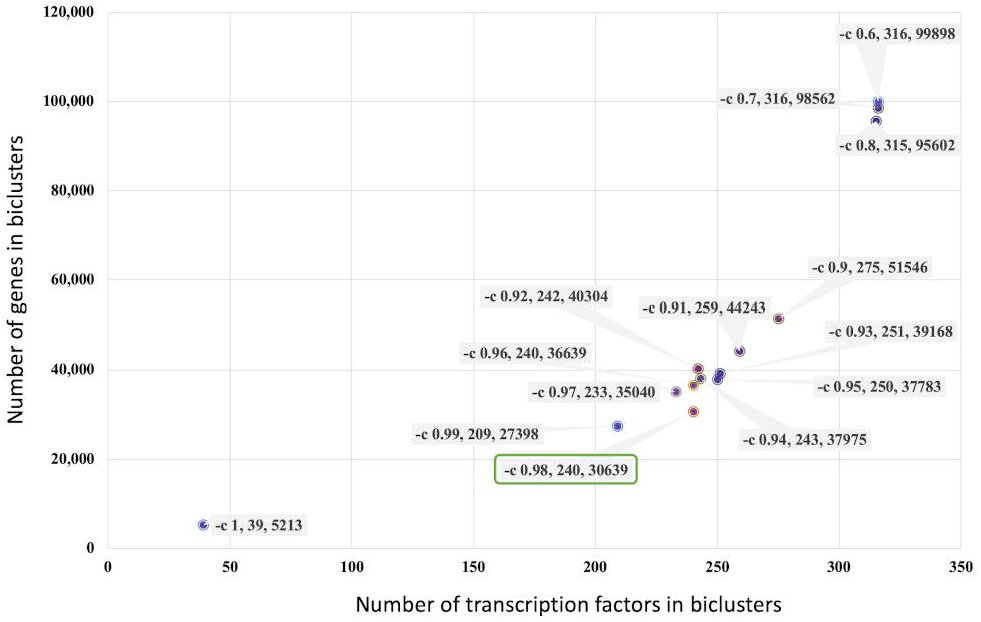
Optimization of QUBIC parameter. Relationship between QUBIC’s consistency parameter “*c*” (tested from 1 to 0.6) and the number of target transcription factors (TFs) included in bicluster (BC) versus the size of the BC (total number of genes). Each block denotes –c value, TF included in BCs, and total number of genes at that “*c*” value. The optimal “*c*” value selected for final analysis is highlighted.

### Evaluation of QUBIC biclusters using characterized TFs and co-expressed genes from public lateral root organ-related datasets

We observed organ-specific bicluster each for lateral root (both ELR and LR; BC001) and nodule (both EN and MN; BC013) tissues that included all three biological replicate samples in one bicluster, suggesting that these are likely to be highly consistent and reproducible. Four BCs each were specific to all three replicates of ELR (BC015, 019, 033 and 101) and MN (044, 048, 152 and 155) tissue types (Supplementary Table S3). To test the rationality of BCs, we compared the expression patterns of co-expressed genes with marker TFs in publicly available transcriptome data (19). The transcription factor “*NSP*1 (*Glyma16g01020*)” crucial for nodule development was present in BC037 and BC045 (Supplementary Table S3B). BC037 was specific to nodule tissues and comprised of 367 co-expressed genes. Among these, 52% had more than two-fold up-regulation in EN and MN tissues in our RNA-seq data. A marker gene highly enriched in nodule tissues, *ENOD*40 (*Glyma02g04180*), was found in five BCs (BC013, 22, 40, 45, 53 and 95) with different combinations of nodule samples clustered together in each BC. All genes in BC013 that showed specificity for nodule tissue samples with all three replicates in EN and MN in our study. Also, 50% of the genes from this BC showed greater expression in nodule tissue relative to other tissues types in the soybean gene expression atlas (19) (Supplementary Table S4). Gene Ontology (GO) enrichment analysis for this BC showed enrichment of nucleic acid metabolic process GO term with a significant p-value (FDR; 0.02) and molecular function GO term “Purine ribonucleoside triphosphate binding (FDR; 0.05); both of which are associated with biological nitrogen fixation, a process specific to nodule tissues. For example, soybean nodules export nitrogen in the form of ureides (purines) (28). The above observations indicate the appropriate clustering of relevant transcripts and validate the parameters used for clustering. Notably, we observed few novel transcripts and genes with unknown function, co-expressed in the nodule-specific biclusters (Supplementary Table S4). This observation suggests a potential role for these genes in nodule development and offers candidate genes for functional characterization.

Further, we took advantage of the time course data for IAA-induced lateral root development in *Arabidopsis* (29), to select and evaluate marker genes present in LR-related BCs in soybean. For example, the LR marker TF, *GmTMO7* (*Glyma04g34080*), a potential ortholog of Arabidopsis *TMO7* identified in the above study, was present in BCs 110, 120 and 173 (Supplementary Table S3). Of the 113 genes present in BC120, 96 showed coordinated up-regulation with *TMO*7 in LR tissues, whereas 17 showed negative co-expression. Upon comparison with the Arabidopsis LR induction time course dataset (29), we found 15 co-expressed soybean orthologs (13 positively co-expressed and 2 negatively co-expressed). Where seven (out of 13) from positively co-expressed gene orthologs set were mostly induced in the later stage of lateral root development, one (out of two) from negatively co-expressed had down-regulation in a later stage of lateral root development (see marked blue and red box in Supplementary Table S5). The other lateral root marker *LRP*1 was in BC019 that comprised of 845 genes. Among these genes, 746 were positively and 99 were negatively co-expressed with LRP1 in all three replicates of ELR. Interestingly, 30 (out of 46 matched genes) of the positively co-expressed genes were potential orthologs of Arabidopsis genes that also showed induction during a similar stage of lateral root development (Supplementary Table S5) in the LR induction time course dataset (29). These comparisons enabled us to evaluate the ability of biclustering parameters and GRN algorithms to appropriately identify regulators and regulatory relationships of target genes during root lateral organ development.

### Regulatory TF and their Gene Regulatory Networks (GRN) related to root lateral organ development in soybean

For the prediction of regulators and inference of corresponding GRNs, we utilized only those 141 BCs that contained our TFs of interest and marker TFs (240 TFs) which comprised 25.8% (29,270 out of 113,210) of expressed gene transcripts. This approach potentially reduced the computational complexity and time required for modeling GRNs relevant to our study. This sum expression matrix of 29,270 genes and 240 TF genes (Supplementary Table S6) was used as input for GRN inference by Lemon-Tree mode I, mode II and Inferelator.

Lemon-Tree produced 828 tight clusters in step 1 from the input expression matrix. A higher number of clusters (828 vs. 141 BCs from QUBIC) suggested that Lemon-Tree clusters were relatively more discrete/smaller in comparison to QUBIC BCs. In step 2, two separate options/modes were utilized (See methods and Figure 1). In mode I, we utilized the 828 tight-clustered modules generated by Lemon-Tree (mode I) and in mode II, the 141 BCs produced by QUBIC (mode II). In mode I, 176 TFs were ranked as the top 1% regulators, whereas in mode II, 92 TFs were ranked as top 1% regulators (Supplementary Table S7). Score evaluation was performed for top 1% and randomly predicted regulators from both modes. In both the cases, the minimum score for a top regulator (14.22; mode I and 12.13 mode II) was ~3 times higher than the maximal score (4.99; mode I and 4.23; mode II) for a randomly assigned regulator (Figure 4). This suggested that the scores for top regulators are greater than what could be expected by chance. Inferelator algorithm predicted 132 TFs as potential regulators and five predicted groups (Supplementary Table S7). Comparison of 176, 92, and 132 TFs predicted as regulators respectively, by Lemon-Tree mode I, mode II, and Inferelator, revealed that 56 TFs (~27%) were predicted by all three different modes (Figure 5A). We ranked these common 56 TFs as high confidence TF regulators. In addition, ~62% of the TFs predicted as regulator by Lemon-Tree mode I were also identified as regulators by Lemon-tree mode II and/or Inferelator (Supplementary Figure S1A).

**Figure 4.**
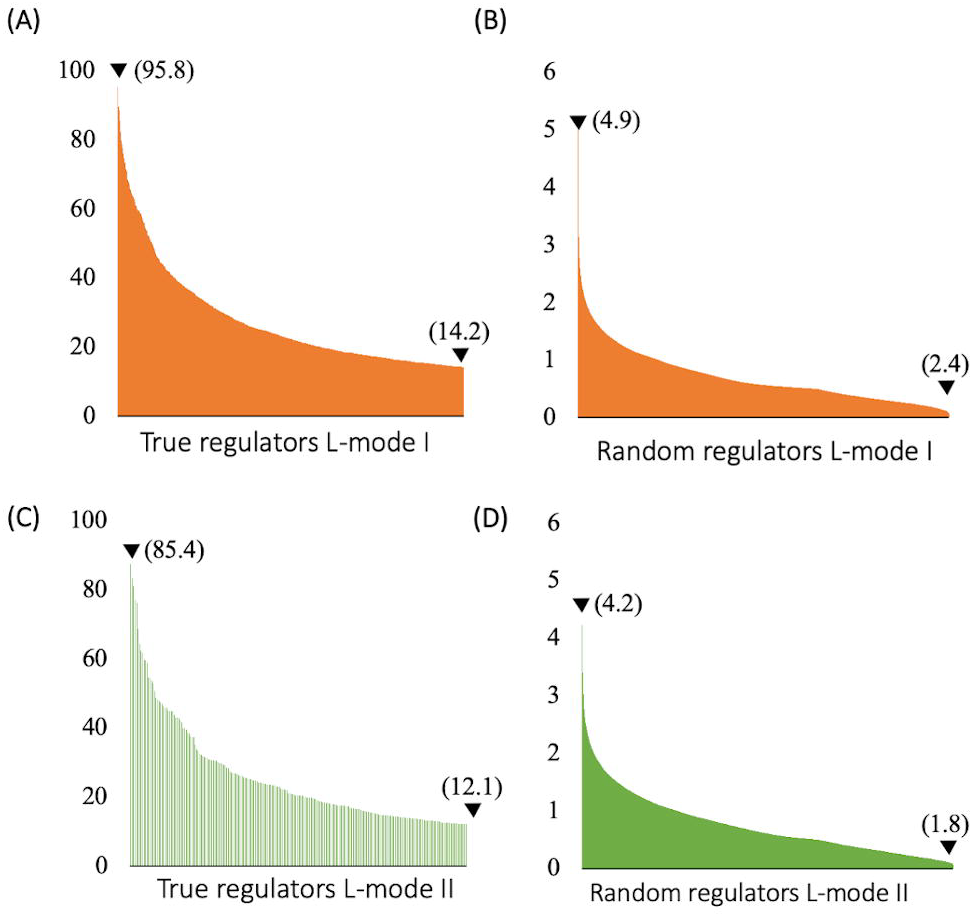
Distribution of Lemon-Tree scores of true and random regulators for root lateral organ development in soybean. Histogram shows the distribution of score for true and randomly assigned regulator from Lemon-Tree mode I (orange) and mode II (green) produced network. Arrows indicate the minimum and maximum scores from each category with values in parenthesis.

**Figure 5.**
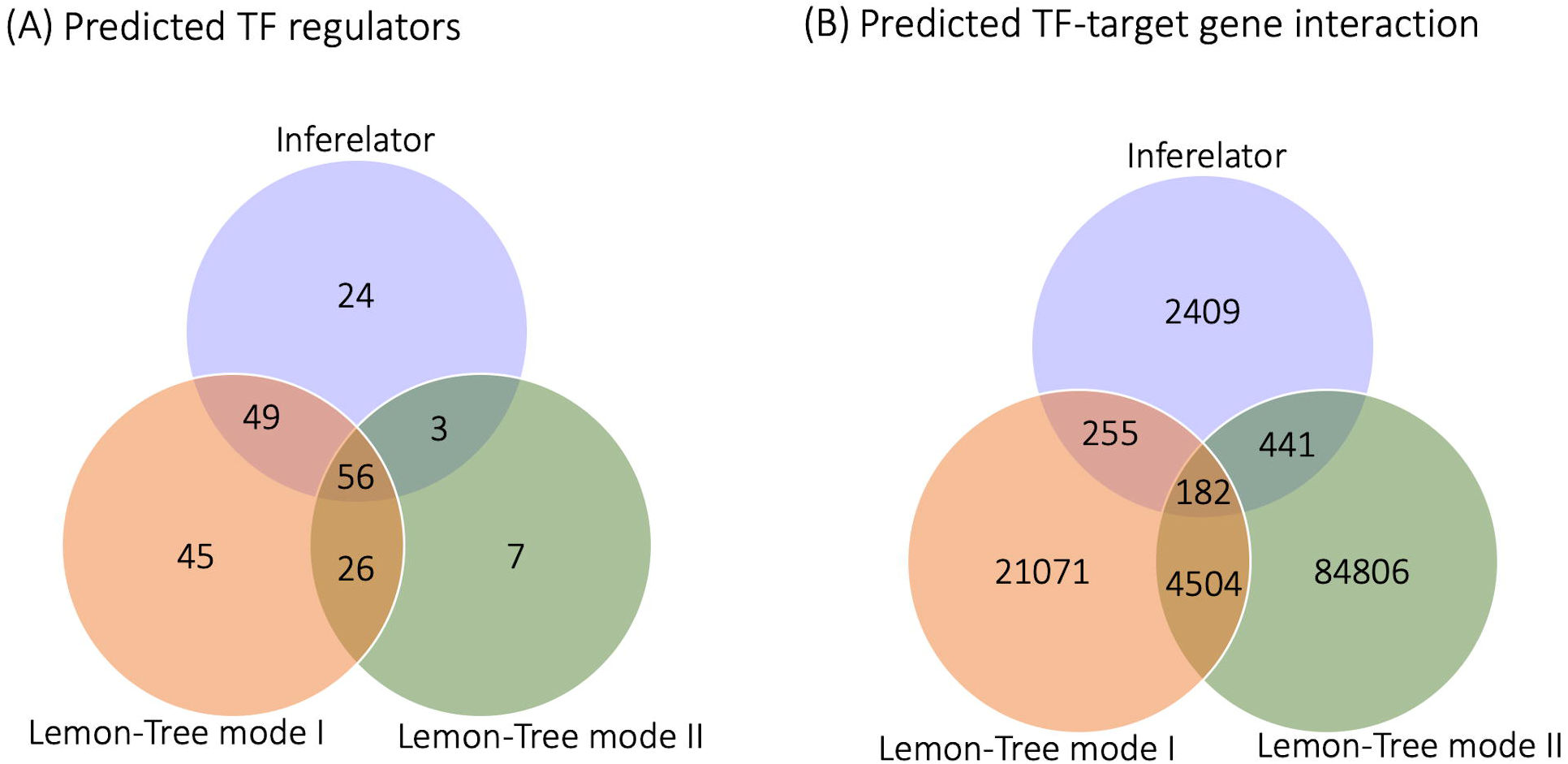
Overlap and differences among outputs from the three different network approaches. (A) Transcription factors (TF) predicted as regulators from three different network approaches. (B) Regulatory interactions predicted by three different network approaches. Numbers in center indicate the number of potential regulators (in A) and interactions (in B) recovered by all the three different approaches playing role in root lateral organ development in soybean.

Furthermore, a total of 113,668 non-redundant TF-target regulatory interactions were predicted by all three modes (Lemon-Tree mode I – 26,012, mode II – 95,845 and Inferelator - 3,287) (Supplementary Table S8). A higher number of regulatory interactions in Lemon-Tree mode II is likely due to larger BCs produced by QUBIC. There was relatively smaller overlap among the three modes (Supplementary Figure S1B). We evaluated if the known LR and nodule marker TFs were predicted as regulators as a measure of successful TF prediction by the three different modes. Soybean orthologs of lateral root marker TFs, LRP1 (Glyma14g03900), ARF5 (Glyma14g40540), CRF2 (Glyma08g02460), and *TMO7* (Glyma04g34080 and Glyma06g20400) were predicted as regulators by all three inference modes. Additional orthologs of ARF5 (Glyma17g37580) and CRF2 (Glyma05g37120) were predicted as regulators by Lemon-Tree mode I and II. However, orthologs of GATA23 (Glyma03g39220, Glyma19g41780), and LRP1 (Glyma02g44860, Glyma07g35780) were not identified as regulators by any of the modes. These four genes were not enriched in LR tissues (Supplementary Table S2) potentially why they were not predicted as a regulator in this dataset. Successful prediction of four of the five LR-associated markers correctly as regulators by all three modes suggested that the pipeline was reliable and would be used in predicting previously unknown regulators of nodule development.

A number of TFs were demonstrated to play a crucial role in nodule development through genetic evidence from model legumes (4, 30). These include *NODULE INCEPTION* (*NIN*) (RWP-RK family; (31), NODULATION SIGNALING PATHWAY1 and 2 (NSP1 and NSP2; GRAS domain proteins), Nuclear Factor Y (NF-YA1; (32)), <u>E</u>thylene Response Factors <u>R</u>equired for <u>N</u>odulation (ERN1 and ERN2; AP2/ERF family; (33)), and CYCLOPS (coiled-coil domain protein) (34–37). In addition, a MYB TF that interacts with NSP2, an ARID domain protein that interacts with SymRK, a bHLH and a set of HD-ZIP IIIs involved in nodule vascular development, and a C2H2 Zn finger TF involved in bacteroid development are also known (38). A potential soybean ortholog of NIN, Glyma02g48080 (34), belonging to orthogroup OGEF1237 was predicted as a regulator by Lemon-Tree mode I. Only one other NIN-like gene in this orthogroup (Glyma04g00210) was included in our list of input TFs based on expression enrichment in nodules, but was not predicted as a regulator by any mode. Two other NIN-like genes outside of this orthogroup (Glyma12g05390 and Glyma01g36360) were predicted to be regulators by Lemon-Tree modes I and II. Nodule-enriched NFY-As (Glyma02g35190 and Glyma10g10240) were identified as regulators by Lemon-Tree mode I and Inferelator. In *Lotus japonicus*, two Nuclear Factor-Y (NF-Y) subunit genes, *LjNF-YA1* and *LjNF-YB1*, were identified as transcriptional targets of NIN (39). In agreement, our analysis predicted that one of the soybean NIN-like genes, Glyma12g05390, regulates NF-YA1 (Glyma10g10240; Lemon-Tree mode II) and the other NIN-like gene, Glyma01g36360, regulates NF-YA2 (Glyma02g35190; Lemon-Tree mode I; Supplementary Table S7).

Two potential orthologs of LjERN1 (Glyma02g08020 and Glyma19g29000) were predicted as regulators by Lemon-Tree modes I and II. Among the major nodulation TFs, only NSP1 was not predicted to be a regulator by our GRN pipeline. In summary, the pipeline correctly predicted known nodulation and LR TFs including the expected relationships between NIN, NF-YA, and ERN1.

### Putative protein-protein interactions (PPI) identified in root lateral organ-related GRNs

Co-expressed and co-regulated genes have a higher likelihood of having an indirect functional interaction or direct physical interaction (40). Many TFs form a complex with other proteins for proper molecular and cellular activity. PPIs are the physical interactions between two or more proteins which form the crux of a functional protein complex formation (41). To evaluate if potential regulators identified by us undergo PPIs with other co-regulated proteins, we compared all 113,668 unique TF-target predicted regulatory interactions from three modes of GRN inference method against experimentally verified and/or predicted PPIs based on experimental data reported in the STRING database (see methods for details). We identified, 843 potential interactions among 69 TFs with PPI confidence scores ranging from 150 to 995 (Supplementary Figure S2, Supplementary Table S9). The high scorer (>800) PPIs were observed from Lemon-Tree mode II run. It was previously suggested that a score < 800 were probably false positives that originated from prediction methods (42). Also, the maximum number (~64%) of PPI interactions were identified by Lemon-Tree mode II, while only four PPI were predicted by all three modes (Supplementary Figure S1C). A likely explanation is the comparatively bigger BCs in this mode generated by QUBIC. While overall, in comparison to all predicted interactions by each mode independently, Inferelator had a greater frequency (2%) of interactions in PPI, i.e., out of total predicted 3288, 61 were observed in PPI, followed by Lemon-Tree Mode I (1%) and then mode II (0.65%). Two ARF5 lateral root markers Glyma14g40540 and Glyma17g37580 were predicted to interact with Glyma13g43050 (PPI score 980) and Glyma15g13640 (PPI score 530) present in GRNs predicted by Lemon-Tree mode I and Inferelator respectively. Glyma13g43050 is an ortholog of Arabidopsis IAA28 which has been demonstrated to interact with AtARF5 (43), and this regulatory module plays a key role in lateral root development (44).

### High confidence TF regulators and their GRNs associated with root lateral organ development in soybean

To determine high-confident regulatory interactions and build a consensus GRN, we evaluated if interactions were conserved across all three modes of GRN prediction (Lemon-Tree modes I, II and Inferelator). Results showed that 182 co-regulatory interactions (for 21 TFs) were commonly predicted by all three modes (Figure 4B, Supplementary Table S10). Therefore, for 38% of the TFs predicted as a regulator (21 of 56), have also predicted common target genes independently by all three modes. These 21 TFs made independent GRN with their co-regulated target genes (Figure 6). We ranked the consensus interactions by computing the average of the normalized score given by all three GRN inference modes (ranged from min = 0.19, max=0.88) (See materials and methods for full detail). Table 1 shows the score for 21 commons TFs and their common regulatory interaction predicted from different methods (Lemon-Tree mode I, Lemon-Tree mode II and Inferelator). The complete list of modules together with their high-scorer regulators for this study is available in the Supplementary Table S10. Based on the expression of the TF regulator and their predicted target (Figure 7), we categorized GRN enriched in specific lateral organ tissues.

**Table1.**
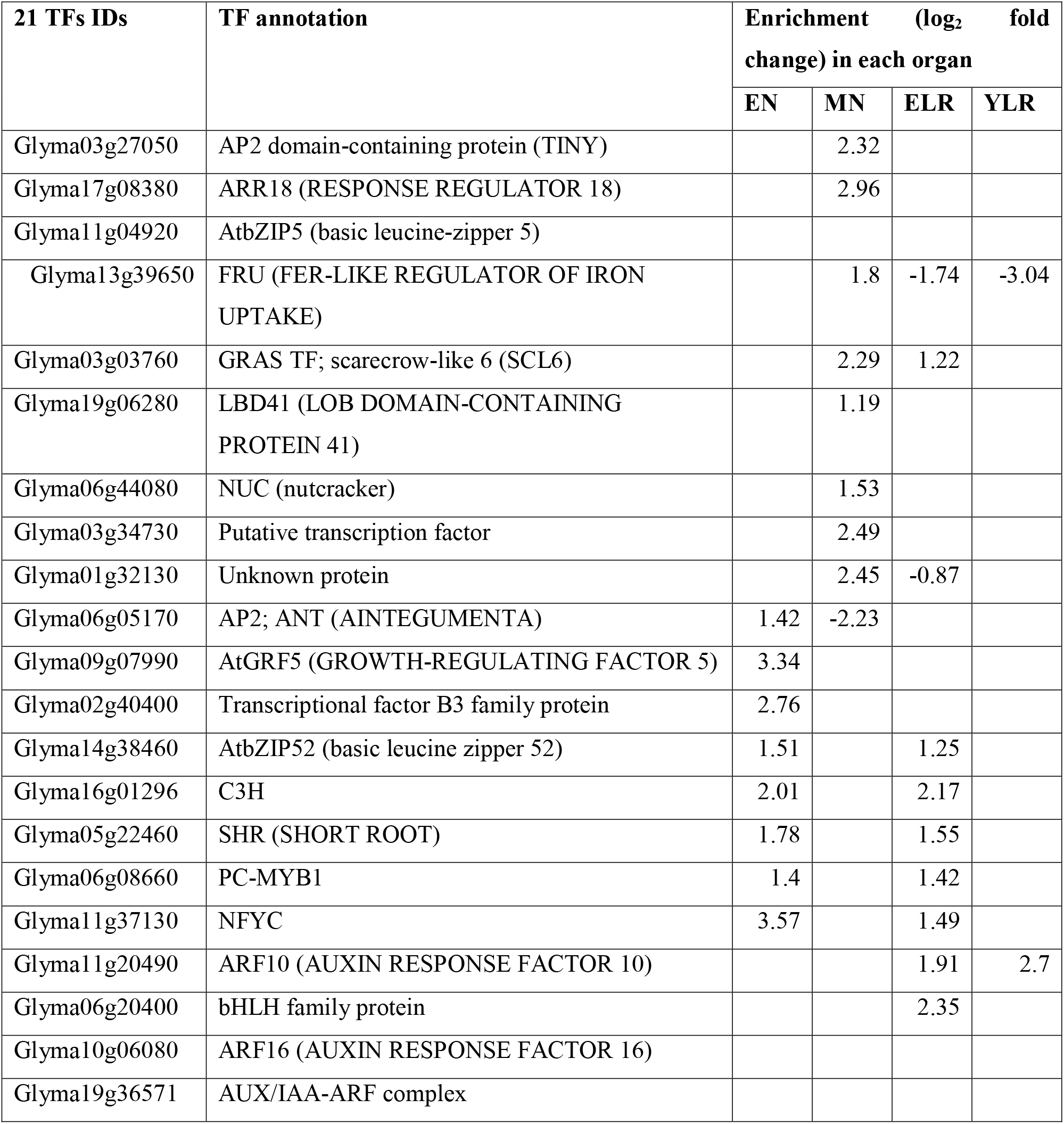
List of transcription factors predicted as regulator by all three workflows used in our study.

| 21 TFs IDs | TF annotation | Enrichment (log<sub>2</sub> fold change) in each organ |  |  |  |
| --- | --- | --- | --- | --- | --- |
|  |  | EN | MN | ELR | YLR |
| Glyma03g27050 | AP2 domain-containing protein (TINY) |  | 2.32 |  |  |
| Glyma17g08380 | ARR18 (RESPONSE REGULATOR 18) |  | 2.96 |  |  |
| Glyma11g04920 | AtbZIP5 (basic leucine-zipper 5) |  |  |  |  |
| Glyma13g39650 | FRU (FER-LIKE REGULATOR OF IRON UPTAKE) |  | 1.8 | -1.74 | -3.04 |
| Glyma03g03760 | GRAS TF; scarecrow-like 6 (SCL6) |  | 2.29 | 1.22 |  |
| Glyma19g06280 | LBD41 (LOB DOMAIN-CONTAINING PROTEIN 41) |  | 1.19 |  |  |
| Glyma06g44080 | NUC (nutcracker) |  | 1.53 |  |  |
| Glyma03g34730 | Putative transcription factor |  | 2.49 |  |  |
| Glyma01g32130 | Unknown protein |  | 2.45 | -0.87 |  |
| Glyma06g05170 | AP2; ANT (AINTEGUMENTA) | 1.42 | -2.23 |  |  |
| Glyma09g07990 | AtGRF5 (GROWTH-REGULATING FACTOR 5) | 3.34 |  |  |  |
| Glyma02g40400 | Transcriptional factor B3 family protein | 2.76 |  |  |  |
| Glyma14g38460 | AtbZIP52 (basic leucine zipper 52) | 1.51 |  | 1.25 |  |
| Glyma16g01296 | C3H | 2.01 |  | 2.17 |  |
| Glyma05g22460 | SHR (SHORT ROOT) | 1.78 |  | 1.55 |  |
| Glyma06g08660 | PC-MYB1 | 1.4 |  | 1.42 |  |
| Glyma11g37130 | NFYC | 3.57 |  | 1.49 |  |
| Glyma11g20490 | ARF10 (AUXIN RESPONSE FACTOR 10) |  |  | 1.91 | 2.7 |
| Glyma06g20400 | bHLH family protein |  |  | 2.35 |  |
| Glyma10g06080 | ARF16 (AUXIN RESPONSE FACTOR 16) |  |  |  |  |
| Glyma19g36571 | AUX/IAA-ARF complex |  |  |  |  |

**Figure 6.**
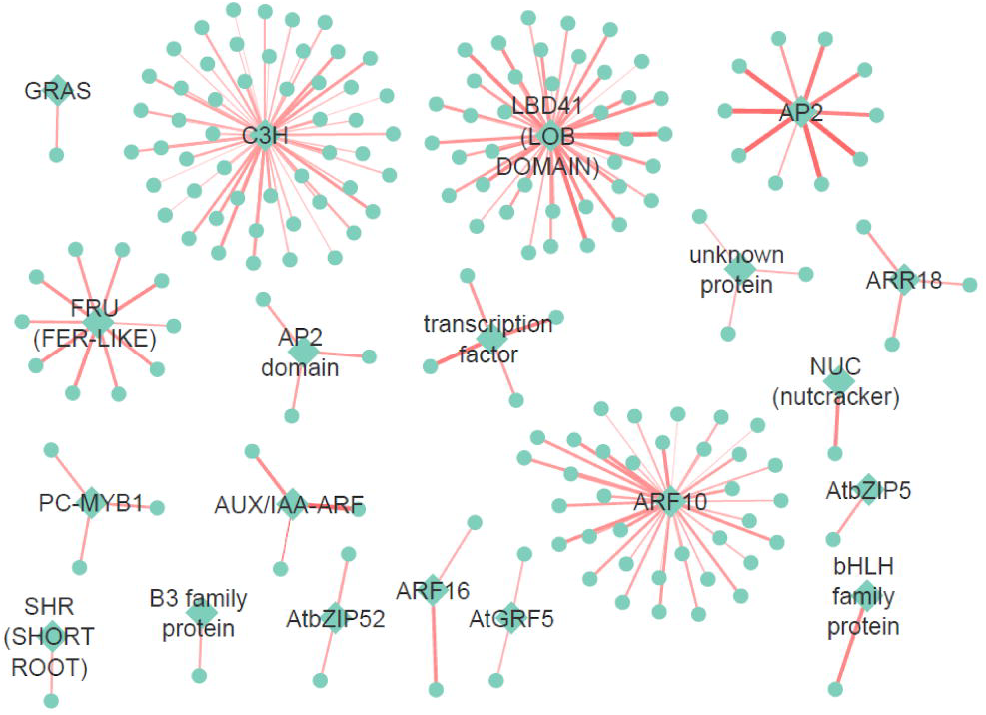
Figure showing consensus 182 co-regulatory interactions predicted and recovered by three different modes chosen in this study. Nodes in diamond denote regulator transcription factors (TFs) and circles denote predicted target genes. Edges denote the normalized score of interaction calculated by all three different modes. Broader the edges, higher the interaction score.

**Figure 7.**
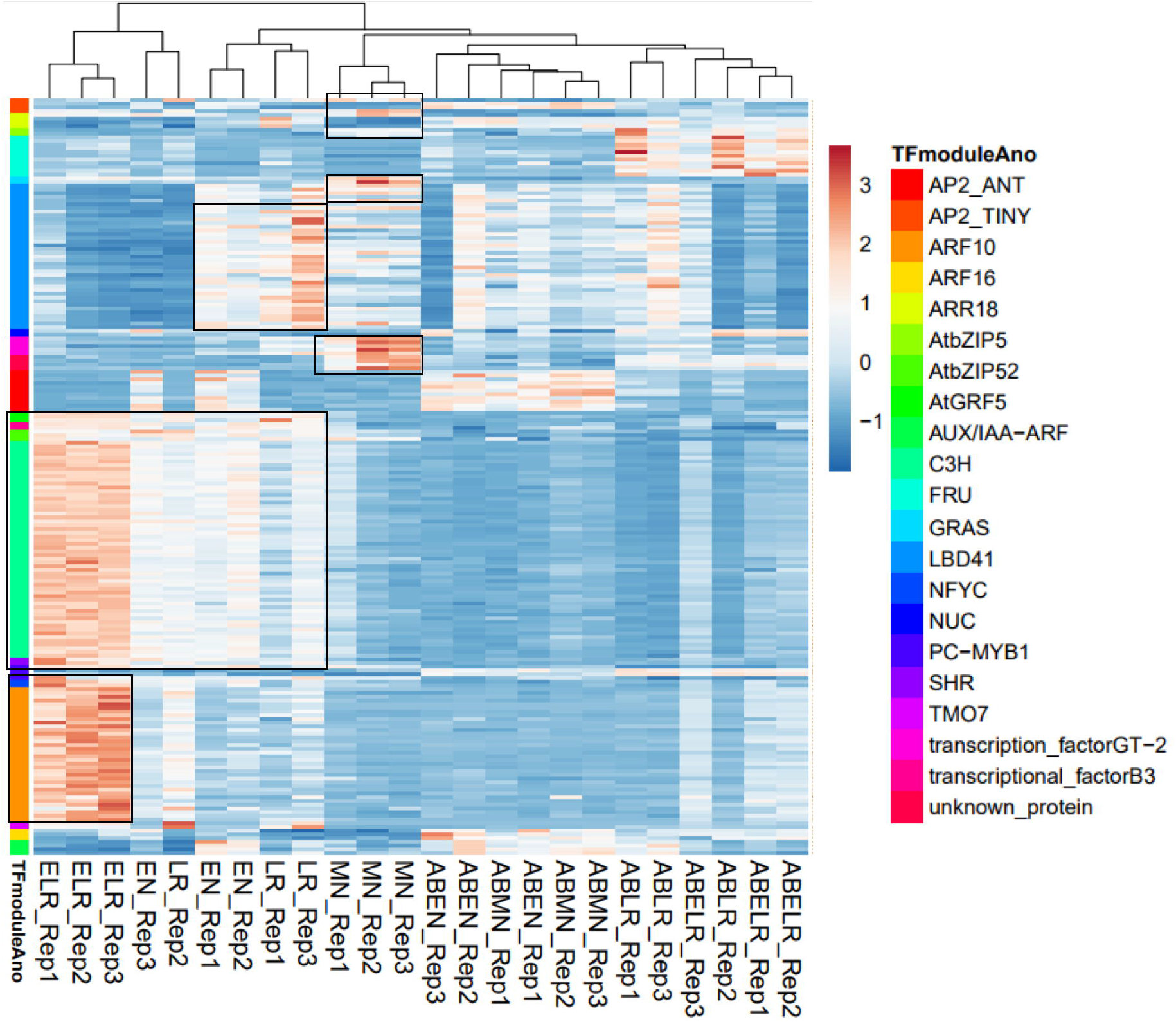
Heat map showing normalized expression from varied samples of root lateral organ development in soybean for regulator transcription factors (TFs) and their co-regulatory target genes in consensus network predicted by three different modes chosen in our study. Row annotation for 21 regulator TFs and their co-regulatory partners are shown in different colors.

TF regulators AP2; ANT (AINTEGUMENTA), transcriptional factor B3 family protein, AtGRF5 (Growth-Regulating Factor 5), C3H, AtbZIP52 (*Arabidopsis thaliana* basic leucine zipper 52), PC-MYB1, and SHR (Short Root) appear to co-regulate GRN modules during early nodule (EN) development. TF regulators GRAS; scarecrow-like transcription factor 6 (SCL6), LBD41 (LOB Domain-Containing Protein 41), AP2 domain-containing transcription factor TINY, NUC (nutcracker); nucleic acid binding, AtbZIP5 (Arabidopsis thaliana basic leucine-zipper 5), FRU (FER-Like Regulator Of Iron Uptake), ARR18 (Arabidopsis Response Regulator 18) and two unknown TF proteins appear to co-regulate GRN modules late during nodule (MN) development. Interestingly, four PPI interaction (out of total 843 PPI network) were also commonly predicted by all three GRN inference networks in our study for LBD41 and FRU in mature nodules (Supplementary Table S10, Supplementary Figure S2B). ARF16 and AUX/IAA-ARF complex were observed for ELR development, whereas TMO7 and ARF10 (Auxin Response Factor 10) co-regulated GRN for YLR development in soybean.

## Discussion

In spite of the economic and environmental importance of biological nitrogen fixation in nodule in soybean, there is still an unanswered question of what key TFs regulate the underlying GRNs in nodules and lateral roots (4). We developed a robust computational framework for GRN construction using genome-scale gene expression data. Specifically, this framework integrates genomic and transcriptomic data to infer the key regulators and GRN associated with nodule development in soybean. The predicted networks consistently included experimentally verified genes, demonstrating the ability of our framework to reveal significant, potentially important GRNs. With a broader impact, the framework can be used as a template for constructing GRNs to address any biological question of interest in any species.

To reduce the computational complexity and make the predicted regulator TFs and GRNs relevant to our biological question, a biclustering method and a regulatory network inference tool were used, where their parameters were optimized via several iterations for data analysis and modeling. Among existing GRN inference algorithms, Lemon-Tree and Inferelator were successfully applied in different biological questions due to their valued feature i.e. top regulator and top-ranked regulatory target prediction (45–48). Lemon-Tree detects regulatory modules and regulators from gene expression data using probabilistic graphical models (17). Whereas, Inferelator learns a system of ordinary differential equations using the Bayesian Best Subset Regression that describes the rate of change in transcription of each gene or gene-cluster, as a function of TFs. It has been shown that predictions made by the Inferelator are highly accurate for top ranking predictions. Stochastic Lemon-Tree and Inferelator perform better if the transcriptional program can be inferred from a pre-specified list of regulators rather than from a full gene list, because erroneous interactions with non-regulators will be eliminated a priori (49). So, we took the differentially expressed TFs and predefined marker TFs with a known role in nodule and LRs to infer GRN.

### Novel regulators of nodule development

We distinguished organ (lateral root/ nodule) and/or developmental stage-specific (early/mature) consensus GRNs based on organ-specific enrichment of the TFs, their differential expression and expression pattern of their co-regulated genes in our transcriptome data. In addition, we also employed comparative genomics and information from public tissue atlas and transcriptome data. The analysis correctly predicted four of the five LR regulators with high confidence and known nodulation TFs including the expected relationships between them. For example, the phylogenetic analysis suggested that ERN2 may not be present in legumes that form determinate nodules such as soybean, *L. japonicus*, or common bean (50). The expression of *ERN1* and *ERN2* are under the control of NIN and NF-YA in *Medicago*, a legume that forms indeterminate nodules. In fact, NF-YA binds the promoter of *ERN1* directly regulating its expression in *Medicago*. However, *ERN1* expression does not appear to be regulated by NIN or NF-YA in *L. japonicus* as its expression is not altered in *nin* or *nf-ya* loss of function mutants. Our GRN prediction also did not identify *ERN1* as a target of NF-YA or NIN in soybean. *ERN1* is directly regulated by CYCLOPS in *L. japonicus*. NSP2 and CYCLOPS were not included in the input TF list due to no nodule-specific enrichment and/or incorrect annotation. The inclusion of CYCLOPS in future analyses might reveal regulatory relationships between ERN1 and CYCLOPS in soybean. It remains to be seen if this is conserved among other determinate nodule forming legumes including soybean. Given the reliability of the pipeline in accurately predicting known TFs, we discuss previously unknown regulators of nodule development predicted by the pipeline.

An identified EN-GRN was enriched with cell division and cycle functions. Three TFs were predicted to drive GRNs specifically associated with emerging nodules, which are soybean orthologues of Arabidopsis ANT (AINTEGUMENTA; At4g37750), AP2/B3 domain transcriptional factor (At5g58280), and AtGRF5 (Growth-Regulating Factor 5). All the three genes are associated with sites of cell proliferation in Arabidopsis. While GRF5 plays a role in cell proliferation during leaf primordia formation and leaf development, ANT is crucial for flower development. At5g58280 shows the highest expression level in the shoot apex, particularly in the central zone. Indeed, it is likely that the soybean TFs associated with EN GRNs direct cell proliferation during early nodule development. Seven other TFs belonging to C3H, bZIP, MYB1, NF-YC, and SHR were also predicted to co-regulate GRN modules in both emerging nodules and emerging lateral roots (Table 1). Soybean ANT ortholog was the regulator with the highest score in our analysis (0.8) and was predicted to co-regulate ten target genes specifically in emerging nodules. Its targets included *ATCSLA*09, *ALDH*2C4, *GCL*1 (*GCR*2-LIKE 1), *AAP*6, and auxin-responsive protein. A maximum of 51 co-regulated target genes were predicted for a C3H TF regulator (enriched in both EN and ELR) by all three modes. Most of the target genes such as glycosyl hydrolase family protein, *CYCA*1;1 (Cyclin A1;1), zinc finger (C3HC4-type RING finger), *CDKB*1, *CMT*3 (chromomethylase 3); DNA (cytosine-5-)-methyltransferase, calmodulin-binding protein-related, *CYC1BAT*; cyclin-dependent protein kinase regulator, mitotic spindle checkpoint protein, putative (*MAD*2), *ATARP*7 (Actin-Related Protein 7); structural constituent of cytoskeleton, kinesin motor protein-related, and *CDC*20.1; signal transducer, were high scoring target genes.

GO enrichment analysis of genes involved in EN and EN-ELR GRNs showed significant enrichment of regulation of a cell cycle, movement of a cell or subcellular component, microtubule-based movement, cell division, and cell cycle biological process. (Supplementary Table S10). This is consistent with biological processes known to occur early during lateral organ development. *Cis-regulatory* motif GACCGTTA was enriched in the EN related GRN regulated by a Myb/SANT TF (Supplementary Table S10).

Similarly, MN-GRN involved in mature nodule development was enriched with meristem initiation and growth. Nine TF regulators belonging to GRAS (scarecrow-like transcription factor 6, SCL6), LBD41 (LOB Domain-Containing Protein 41), AP2 domain-containing transcription factor TINY, NUC (nutcracker); nucleic acid binding, bZIP5 (Arabidopsis thaliana basic leucine-zipper 5), FRU (FER-Like Regulator Of Iron Uptake), RR18 (Arabidopsis Response Regulator 18), a Myb/SANT-like DNA binding protein, and a SCREAM-like protein appear to co-regulate GRN modules late during nodule (MN) development. Among these TFs, LBD41 had the highest score (0.77). LBD41 was predicted to co-regulate 38 target genes, among which *PDC*2 (pyruvate decarboxylase-2) had the highest normalized score (0.7). Other targets included *PSAT*, *SRO*2 (similar to rcd one 2), MEE14 (maternal effect embryo arrest 14), zinc finger (AN1-like), SNF2, trehalose-6-phosphate phosphatase, hypoxia-responsive family protein, bHLH, wound-responsive family protein, and ASP1 (Aspartate Aminotransferase 1) with normalized score > 0.5 (Figure 7). Arabidopsis LBD41 is associated with hypoxia response and multiple targets predicted for the soybean ortholog of LBD41 in MN were also associated with hypoxia (51). Nodule oxygen concentrations are highly regulated to enable the proper functioning of the oxygen-sensitive nitrogenase enzyme complex. It is tempting to suggest that soybean LBD41 might play a role in regulating response to hypoxia in MN. The Arabidopsis orthologs of SCL-6 a key regulator in MN, play a role in shoot branching by regulating axillary bud development (52). We had previously suggested that nodules and shoot axillary meristems require a similar hormone balance during development. It is possible that some developmental pathways such as those regulated by SCL6 are shared between these organs. Similarly, the role of Arabidopsis NUTCRACKER protein required in periclinal cell divisions (53), that of FRU in uptake of iron (54), and RR18 in positive regulating cytokinin activity (55) are all consistent with biological processes observed in MN tissues (56, 57). GO enrichment analysis for MN-GRN genes showed enrichment of specification of axis polarity, adaxial/abaxial axis specification, meristem initiation, meristem growth and regulation of meristem growth (Supplementary Table S10). While these processes are known to occur in mature nodules, TFs associated with these processes had not been identified previously. Genes involved in MN-GRN had significant enrichment (P-value ≤ 0.05 FDR) for *cis*-regulatory motifs GGGCCCAC, ACCG and TGTCGG in their upstream regulatory regions. These are likely to be regulated by TCP, AP2 and B3 TFs respectively (Supplementary Table S10). The study has revealed potential TFs associated with different functions in nodule development.

## Data availability

Gene expression data used to construct gene regulatory networks are available in NCBI Gene Expression Omnibus (GEO), accession number GSE129509. Raw data files are available in NCBI’s Sequence Read Archive (SRA) and can be accessed via links available at the GEO record URL: https://www.ncbi.nlm.nih.gov/geo/query/acc.cgi?acc=GSE129509.

## Acknowledgements

Use of South Dakota State University’s high-performance computing clusters for data analysis and technical support from South Dakota State University’s research information technology team (Dr. Brian Moore) are gratefully acknowledged.

