## Supplementary Figures for "Gene regulatory networks associated with lateral root and nodule development in soybean"

(A – Regulator TFs)

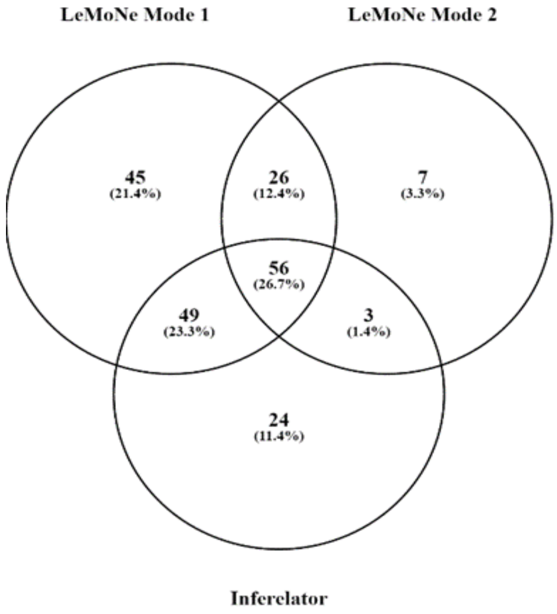


(B – TF-target pairs)

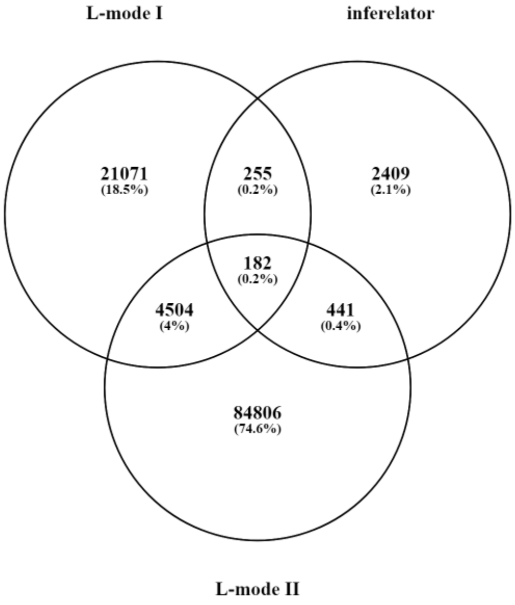


(C – STRING PPIs)

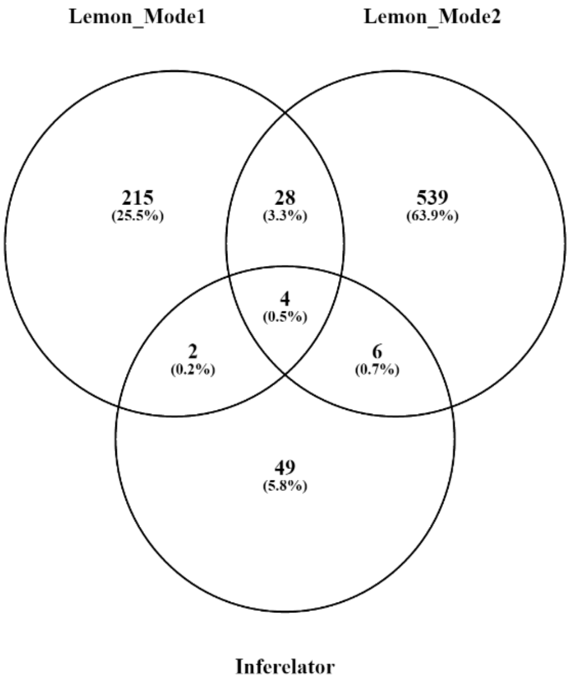


**Supplementary Figure 1**

Venn diagrams outlining overlaps and differences in the outputs among the three different network approaches: (A) Regulator TF prediction; (B) Identification of targets for predicted TFs; (C) Protein-Protein Interactions.

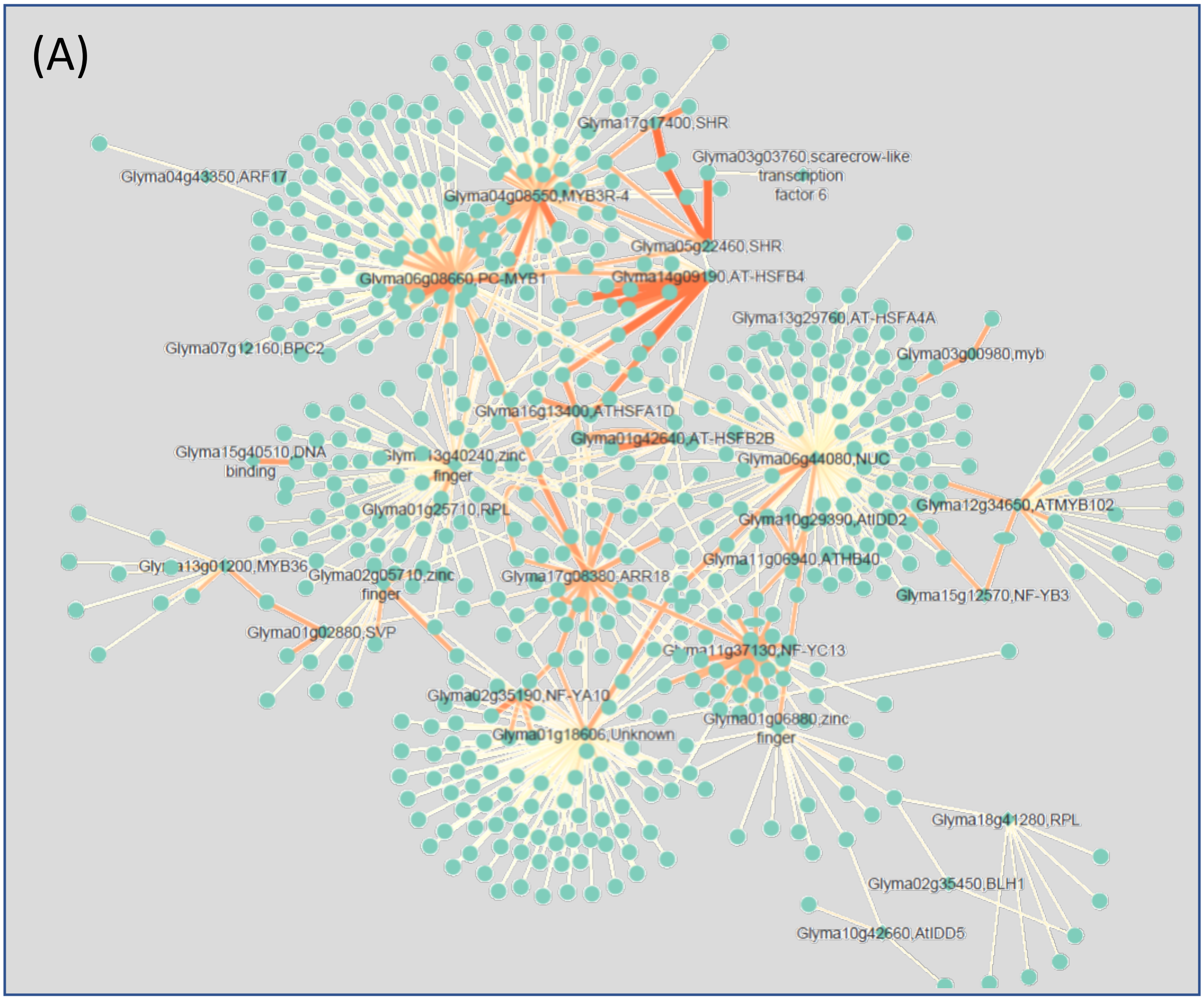


**Supplementary Figure 2A**

A total of 843 STRING protein-protein interaction (PPI) predictions matched our co-regulatory expression prediction. The large network is shown here and smaller discreate networks are shown in Supplementary Figure 2B.

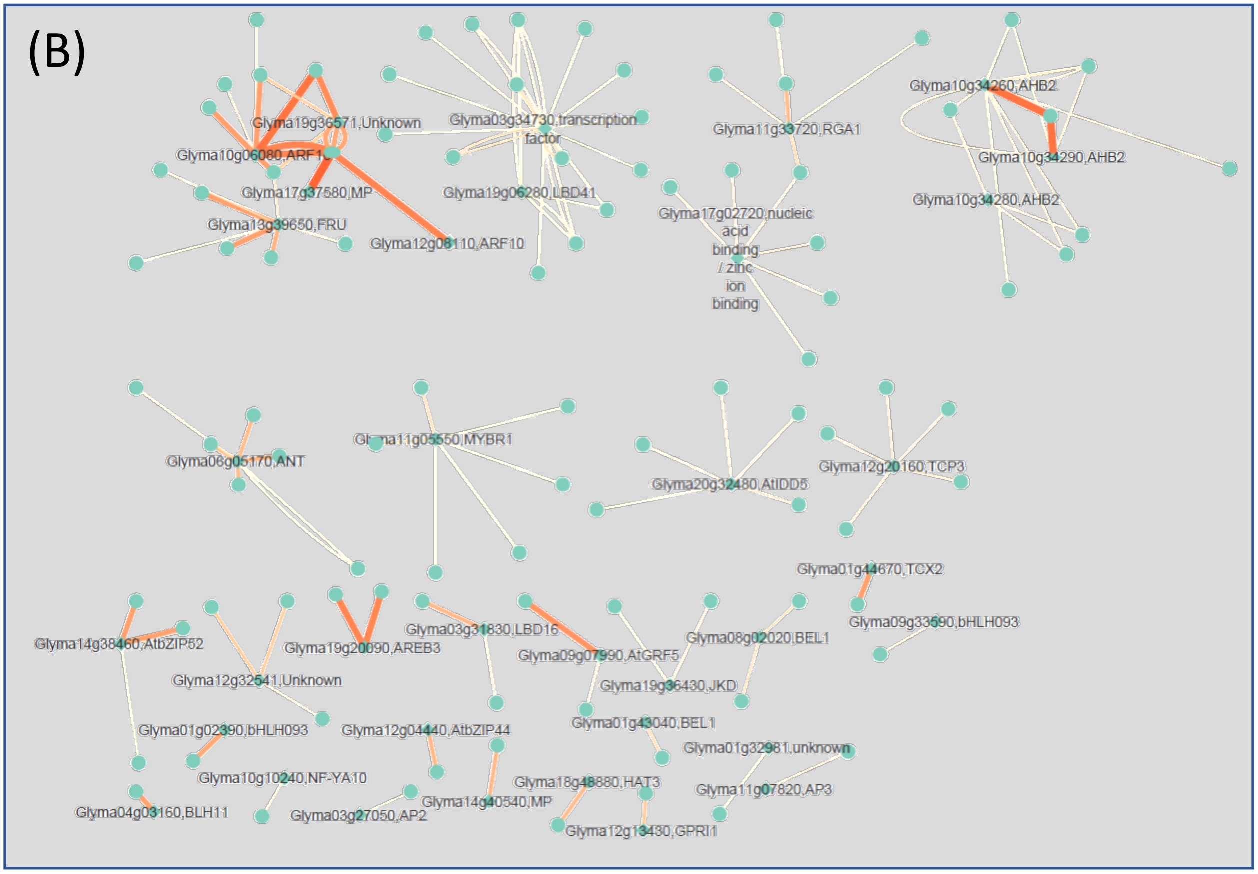


**Supplementary Figure 2B**

A total of 843 STRING protein-protein interaction (PPI) predictions matched our co-regulatory expression prediction. Smaller discreate networks are shown here and the large network is shown in Supplementary Figure 2A.

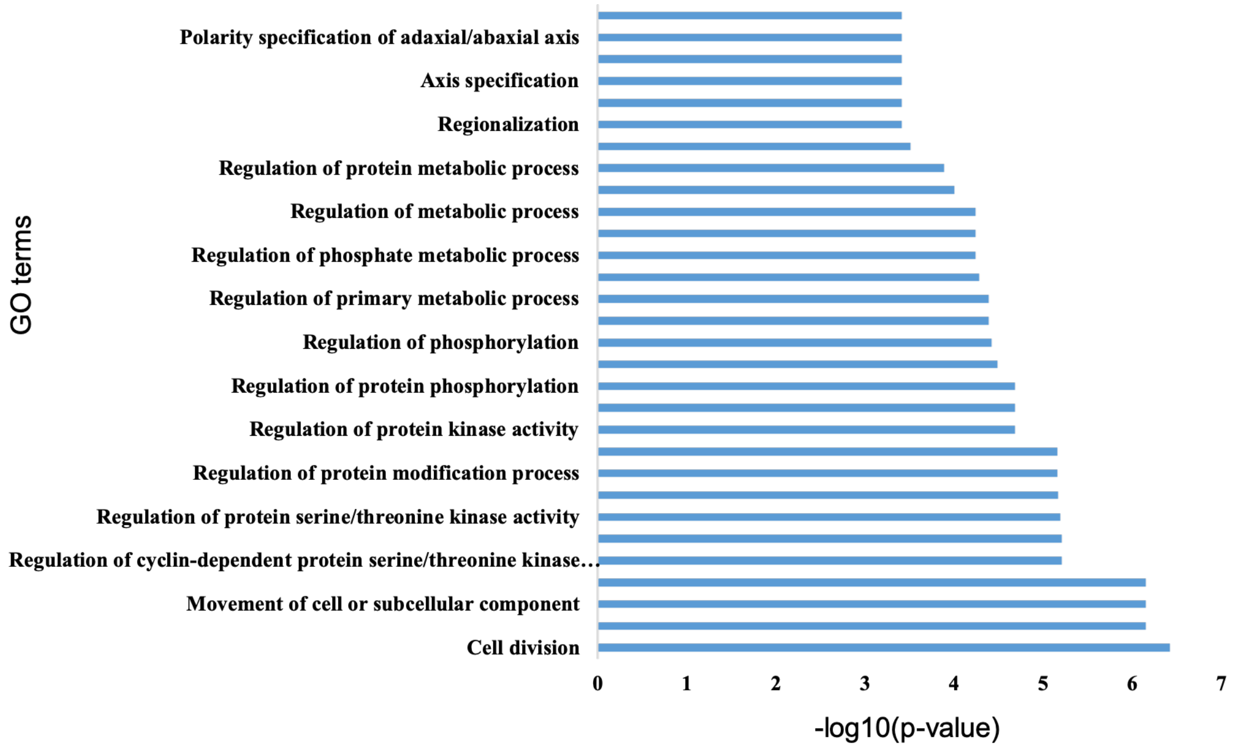


**Supplementary Figure 3**

Gene Ontology biological process enrichment of target genes in consensus 182 co-regulatory gene network predicted for root lateral organ development in soybean.
